# IS-PRM-based peptide targeting informed by long-read sequencing for alternative proteome detection

**DOI:** 10.1101/2024.04.01.587549

**Authors:** Jennifer A. Korchak, Erin D. Jeffery, Saikat Bandyopadhyay, Ben T. Jordan, Micah Lehe, Emily F. Watts, Aidan Fenix, Mathias Wilhelm, Gloria M. Sheynkman

## Abstract

Alternative splicing is a major contributor of transcriptomic complexity, but the extent to which transcript isoforms are translated into stable, functional protein isoforms is unclear. Furthermore, detection of relatively scarce isoform-specific peptides is challenging, with many protein isoforms remaining uncharted due to technical limitations. Recently, a family of advanced targeted MS strategies, termed internal standard parallel reaction monitoring (IS-PRM), have demonstrated multiplexed, sensitive detection of pre-defined peptides of interest. Such approaches have not yet been used to confirm existence of novel peptides. Here, we present a targeted proteogenomic approach that leverages sample-matched long-read RNA sequencing (LR RNAseq) data to predict potential protein isoforms with prior transcript evidence. Predicted tryptic isoform-specific peptides, which are specific to individual gene product isoforms, serve as “triggers” and “targets” in the IS-PRM method, Tomahto. Using the model human stem cell line WTC11, LR RNAseq data were generated and used to inform the generation of synthetic standards for 192 isoform-specific peptides (114 isoforms from 55 genes). These synthetic “trigger” peptides were labeled with super heavy tandem mass tags (TMT) and spiked into TMT-labeled WTC11 tryptic digest, predicted to contain corresponding endogenous “target” peptides. Compared to DDA mode, Tomahto increased detectability of isoforms by 3.6-fold, resulting in the identification of five previously unannotated isoforms. Our method detected protein isoform expression for 43 out of 55 genes corresponding to 54 resolved isoforms. This LR RNA seq-informed Tomahto targeted approach, called LRP-IS-PRM, is a new modality for generating protein-level evidence of alternative isoforms – a critical first step in designing functional studies and eventually clinical assays.

## INTRODUCTION

One of the main applications of mass spectrometry (MS)-based proteomics is characterization of the full complexity of the proteome, including alternatively spliced (AS) protein isoforms.^1–3^ Through alternative splicing, multiple distinct protein spliceforms, or isoforms, can be produced from a single gene. The roughly 20K human genes could give rise to 300K or more protein isoforms, and a rising number of such isoforms have been implicated in diverse processes, ranging from embryonic development^4^ to disease,^5,6,7^ making their direct analytical detection critical.

Despite the prediction of numerous potential proteins, the associated protein sequences are primarily extrapolations from transcript evidence.^8^ Many potential annotated and novel isoforms remain undetected at the protein-level, leaving open questions as to their stability in vivo and their functional roles. Overall, in shotgun MS studies, the peptides that would inform on the presence of particular protein isoforms are detected at extremely low rates.^9,10,3,11^ To illustrate, in one of the most comprehensive MS proteomics efforts to date detected an average of approximately 250 concurrently expressed splice events from trypsin digests per cell line,^12^ despite the potential tens of thousands evidenced by RNA-seq.

Biological and technical issues hinder widespread detection of isoforms via shotgun MS approaches. Alternative isoforms tend to be lower in abundance,^13^ lack uniquely-mapping peptides,^14^ and, through a quirk of evolution, produce fewer proteotypic tryptic peptides on average across splice junctions.^15^ Collectively, these unfavorable properties for AS detection leads to a situation in which valuable isoform-specific peptides are an exceedingly small fraction of a complex mixture, both in identities and quantities. The sampling of such isoform informative peptides is not directed in most shotgun MS frameworks, but rather, is semi-stochastic, wherein DDA or DIA mode is used and the most abundant MS1 precursor peaks are preferentially sampled.^16–19^

In addition to the biological and technical challenges of AS detection, many protein isoforms remain uncharted, entirely missing from reference protein databases.^20^ Many isoforms are still unannotated, uncurated, and the relevant isoform sequence for disease state may not be known.^21^ Incomplete isoform knowledge is underscored by indications from proteogenomic studies^22^.

Recently, our group leveraged new developments in long-read RNA sequencing in which full-length transcripts can be directly sequenced at high nucleotide-level accuracy to experimentally determine transcriptome sequences and their estimated abundances at high depth and coverage.^23,24^ These transcript sequences, being the precursor to protein, are subjected to open reading frame (ORF) prediction and subsequently compiled into full-length protein isoform sequences. Therefore, this “long-read proteogenomics” (LRP) pipeline provides, for a specific sample, the protein isoforms with some prior knowledge of their likelihood of presence *in vivo*. Such isoform sequences could be considered informed hypotheses that require confirmation through appropriate analytical approaches.^25^

For detection of target peptides of interest, targeted MS strategies continue to rapidly improve in their sensitivity and throughput, with increasingly sophisticated downstream computational analysis. Building on foundational targeted MS methodologies, such as selected reaction monitoring (SRM)^26^ and later, parallel reaction monitoring (PRM)^27,28^, a new generation of advanced targeted MS methods are available^29,30,31^ that leverage new capabilities of real-time search^32,17^ machine learning-based prediction of protein features,^33^ and multiplexed isobaric labeling schemes.^34,35^ In recent years, advanced targeted MS approaches have been developed that employ highly multiplexed analysis, allowing for the targeting of hundreds and even thousands of peptides.^29^

One family of advanced targeted MS strategies utilizes dynamic retention-time (RT) adjustment. These methods use information about experimental or predicted peptide elution order to dynamically predict the retention times or target peptides and thus make real-time decisions about MS2 scans, optimizing parameters such as MS duty cycle and, ultimately, sampling sensitivity^36,17,19^. The most sophisticated manifestations of these methods include MaxQuant.Live^37^ and GoDig.^38^

Another family of methods utilizes stable isotope labeled peptides to prompt a “triggering” of MS2 scan collection events directed towards the precursor mass of the endogenous peptide, originally termed Internal Standard-Parallel Reaction Monitoring (IS-PRM) by the Domon laboratory.^39^ SureQuant^40^ and Tomahto^41^ utilize a set of synthetic labeled peptide triggers spiked into the sample of interest. These methods demonstrate highly sensitive, large-scale (i.e. SureQuant ∼600, Tomahto ∼500 target peptides) and multiplexed sampling.

Collectively, these targeted MS approaches demonstrated increased throughput, repeatability, and sensitivity. Assays have been developed on previously detected peptides corresponding to genes in known diseases or pathways, such as tissue specific aging,^41^ Small Ubiquitin-like Modifier (SUMO) and ubiquitin proteomics,^42^ early endosome characterization,^43^ and lipid homeostasis.^38^ Largely, the focus of these studies is the use of advanced targeted MS for improving analytical figures of merit related to quantitative performance.

To address the problem of AS detection, we propose bringing together our recent developments in LRP, which predict alternative protein isoforms, with MS multiplex targeting approaches, thus improving sensitivity to target peptides that are most relevant for isoform discovery and characterization. Our idea is to repurpose advanced targeted MS not for improved quantification of previously detected peptides of interest, as has previously been demonstrated, but, rather, for discovery of undetected or unannotated peptides.^44^ We reason that a trigger-based strategy that employs synthetic peptide spike-ins may increase sensitivity and confidence of AS peptide detection. Sensitivity is derived from the multiplexed targeted scheme, and confidence of novel peptide presence is bolstered by the concurrently fragmented standard peptide and the endogenous peptide within the native matrix and LC/MS run. This schema could provide critical confirmatory support for novel peptides.

In this study, we demonstrate the first application of large-scale targeted detection of peptides predicted from a proteogenomic workflow, which we term long-read proteogenomics IS-PRM, or LRP-IS-PRM. LRP-IS-PRM directs targeted MS methods for purely theoretical, previously unobserved peptides (except within the context of prior transcript data). We describe a human stem cell model system in which we proteogenomically predict alternative isoforms^45^ and target their associated peptides using a recently developed IS-PRM strategy, Tomahto.^41^ Using over one hundred peptide synthetic standards, we demonstrate the benchmarking and application of Tomahto for increased detection of isoform-specific peptides. Lastly, we show how IS-PRM-enabled peptide identifications provide a richer interpretation of protein isoforms through examples of identified peptides placed in the context of a genomic framework.

## METHODS

### Cell Culture

The human iPSC lines used in this study were first generated from a healthy male patient, WTC11^45,46^ using the episomal reprogramming method.^47^ Informed consent was obtained for this procedure. iPSCs were maintained on Matrigel (Corning) in mTeSR Plus media (StemCell Technologies), which was exchanged every other day. For passaging, at ∼70% confluency, iPSCs were passaged using Versene (Gibco) and re-plated in mTeSR Plus + 10 µM Y-27632, a Rho-associated kinase (ROCK) inhibitor (StemCell Technologies), to promote cell survival, and media was changed the next day to mTeSR Plus. For harvesting, iPSCs at ∼70% confluency were washed once with 1x PBS with no Mg^2+^ or Ca^2+^(DPBS, Thermo Fisher Scientific), passaged with Versene, washed two times with 1x DPBS, pelleted, and frozen at -80°C. To ensure equal numbers of cells during cell pelleting, cells were counted using NucleoCounter NC-200 and Via-1 Cassettes (Chemometec). During harvesting, an aliquot of iPSCs was taken to verify pluripotency status of iPSCs. The presence of pluripotency markers Oct3/4 and SSEA-4 were validated using flow cytometry. Each cell pellet contained approximately 10 million harvested cells and was frozen at -80℃.

### Long-read sequencing for defining reference/alternative isoforms in human cell line

#### RNA extraction and sequencing

Total RNA from the WTC11 passage 79 pellet was extracted using the RNeasy Kit (Qiagen). This extracted RNA was analyzed on an Agilent Bioanalyzer to confirm sufficient concentration and quality of RNA for downstream data generation. As described by our group previously,^24^ complementary DNA (cDNA) was synthesized from the extracted RNA, and the Iso-Seq Express Kit SMRT Bell Express Template prep kit 2.0 (Pacific Biosciences) was used on a Sequel II system to obtain long-read sequence information and output Circular Consensus (CCS) reads.

#### Long-read Proteogenomic (LRP) analysis

From the full-length transcriptome for WTC11, we defined the protein isoforms that could be expressed in the cell line. As described previously,^23,24^ an LRP pipeline was used with the help of the workflow framework Nextflow, which enables scalability and reproducibility. Briefly, the CCS reads from long-read sequencing were processed into high fidelity reads using SMRTLink, the primers were removed on the 5’ and 3’ end using “lima,” and Isoseq3 was used to refine, cluster, align, and collapse the reads. Next, SQANTI3^48^ (version 1.3) was run on the output of Iso-Seq3 to classify and assess the quality of transcripts. From the high confidence set of full-length transcript isoforms, we selected the most biologically plausible ORF for each of the Iso-Seq transcripts. First, we identified candidate ORFs (≥ 50 nucleotides) using Coding-Potential Assessment Tool (CPAT)^49^, then we selected the most plausible ORF using the module ‘orf_calling’ based on the following criteria: coding potential, relation of AUG start site to GENCODE reference start sites, and number of AUGs skipped to reach the ORF start site. Transcripts were then grouped by ORFs in the same sequence, and transcript abundance was expressed as full-length read counts per million (CPM). We used SQANTI Protein^23^ and the ‘protein_classification’ module from the LRP pipeline to further classify candidate protein isoforms based on the protein sequence in relation to the reference protein isoforms. Following the nomenclature for transcript isoform classification in SQANTI3, we classified each isoform as one of the following: protein full splice match (pFSM), novel in catalog (pNIC), novel not in catalog (pNNC), or incomplete splice match (pISM). To account for genes that were not detected in the WTC11 long-read (LR) profile, we created a hybrid database by combining the sample-specific FASTA file with known sequences from the GENCODE v38 basic translated transcriptome, hereafter called the “WTC11 protein database.”

### Selection of AS192 peptides

#### AS192 target peptide selection

The LRP-derived WTC11 protein database was subjected to *in silico* tryptic digestion (no missed cleavages, 9-20 amino acids in length). The resulting peptides were filtered to retain ones that were isoform-specific, defined as having an amino acid sequence that maps to only one transcript isoform in the database. Peptides were categorized as being specific for either the major isoform (corresponding to the isoform with the highest transcript abundance) or minor isoform (transcript abundance is less than that of the major gene isoform). A transcript abundance minimum of approximately 10 CPM was selected for the target candidates. Further filtering was done to retain the peptides from genes containing both the major isoform and at least one minor isoform-specific peptide. Of these, a group of 192 peptides was selected (see **Supplemental Table S1**) that correspond to endogenous WTC11 major/minor isoform-specific target peptides.

**Figure 1.**
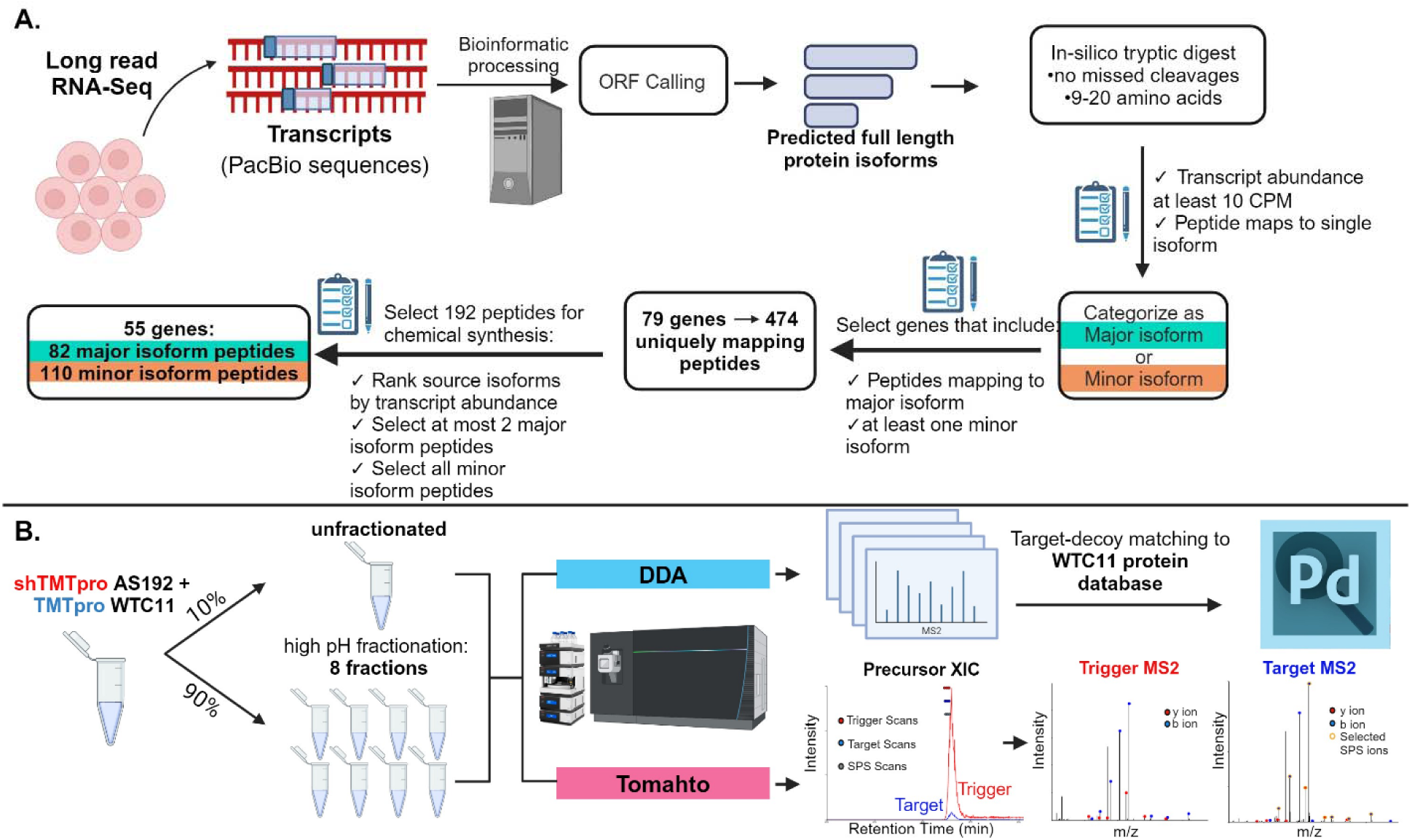
Experimental workflow for isoform-specific peptide selection and targeted MS. A) Selection of isoform-specific peptides inferred from long-read RNA-seq. An LRP bioinformatic workflow was used to predict full-length protein isoform sequences from full-length transcripts detected for a human cell line (WTC11). Target peptides were selected which specifically map to major and minor isoforms of the source gene. A set of 192 peptides were synthesized for MS targeting (AS192). B) Schema of DDA and Tomahto analysis for target peptide detection. WTC11 tryptic peptide digest was labeled with TMTpro and spiked with the pool of 192 synthetic trigger peptides (AS192) labeled with shTMTpro. A 10% aliquot was reserved (unfractionated) while the remaining 90% was subjected to offline high pH fractionation (eight fractions). Tomahto and DDA experiments were performed to detect the target, endogenous AS192 peptides within the WTC11 sample.

### Peptide Standards

#### Peptide synthesis of 192 Isoform-distinguishing target sequences (AS192)

The 192 isoform specific peptide sequences were synthesized by Vivitide (now Biosynth, Gardner, MA, USA) with carbamidomethylation on all cysteines. Peptides were received as crude dry powder with an average of 75% purity and an average yield of ∼1.5 μmol. Peptides were reconstituted in 1000 μL of 20% acetonitrile in water for a concentration of 1.5 nmol/μL. For peptides that did not dissolve completely, 1 μL of concentrated ammonium hydroxide was added. Due to factors, such as synthetic impurity, extent of solubilization, and variability in ionization efficiencies, equimolar mixing of reconstituted peptides yielded a wide range of ion signals for each peptide. To normalize the ion signals so that the ion abundances of most peptides were between 1E7 and 1E8 ion counts, we repooled the synthetics (see **Supplemental Methods**) for unfractionated triggering experiments (see **Results**) with WTC11.

### Proteomics Analysis

#### Sample Preparation

WTC11 cell pellets were lysed with probe sonication using the following lysis buffer: 6% sodium dodecyl sulfate (SDS), 150 mM dithiothreitol (DTT), 75 mM Tris-HCl pH 8. Following lysis, a 660 nm protein quantitation assay (Pierce, ThermoScientific) was performed. Filter-Aided Sample Preparation (FASP) protocol^50^ with 30 kDa, 0.5 mL capacity spin filters (Millipore) for buffer exchange steps was performed, and denaturant buffer (8 M urea in 0.1 M Tris-HCl) was exchanged to final digestion buffer (50 mM ammonium bicarbonate (ABC), pH 8) prior to proteolytic digestion. A 1 μg aliquot of trypsin (Promega) was added to each 100 μg preparation of reduced and alkylated proteins and incubated at 37℃ for 18 h. Peptides were then collected by centrifugation with a final wash of 50 mM ABC. NanoDrop (Thermo Fisher Scientific) analysis at A260 was performed to estimate peptide content.

#### Desalting

Pierce Peptide Desalting Spin Columns (Pierce, ThermoScientific) were used for desalting prior to MS analysis. Samples were ensured to be pH 3 or less, and the manufacturer’s protocol was used with a substitution of 0.1% trifluoroacetic acid (TFA) with 0.1% formic acid (Optima LC/MS grade, Thermo Fisher Scientific).

#### TMT labeling

##### Super-heavy TMTpro (shTMTpro) labeling

Nanodrop was used to assess peptide concentration, and an aliquot equal to 100 µg AS192 peptide or 100 pmol bovine serum albumin (BSA) digest (Pierce, ThermoScientific) was used for (shTMTpro) labeling. Sample was dried via speed vac and reconstituted in 100 µL 0.2 M EPPS buffer, pH 8. An aliquot of 20 µL of MS grade anhydrous acetonitrile (ACROS Organics) was used to reconstitute 0.5 mg of shTMTpro reagent and incubated at room temperature for five minutes with periodic vortexing. The entire 20 µL volume of shTMTpro solution was added to the AS192 or BSA peptide sample. The reaction was incubated for one hour at room temperature, with vortexing every 10 minutes followed by a final centrifugation of the tubes. A 2% aliquot of sample was removed for desalting and was analyzed via LC-MS (TMT labeling quick check method) to check the labeling efficiency. The remaining sample was kept frozen at -80°C until the next day. Samples were treated again with shTMTpro when necessary to achieve 90% labeling efficiency (or sufficient presence of the shTMTpro-labeled peptide as assessed by manual inspection of MS data). For quenching, sample was thawed, and a 5 µL aliquot of 5% hydroxylamine was added to the sample, vortexed, spun down, then incubated for 15 min at room temperature. The sample was then dried via speed vac and reconstituted with 0.1% formic acid in preparation for desalting. After desalting elution, the sample was dried via speed vac and reconstituted with 0.1% formic acid.

##### TMTpro Labeling of Peptides

Nanodrop was used to assess peptide concentration, and an aliquot equal to 100 µg of WTC11 tryptic peptides or 100 pmol BSA digest (Pierce, ThermoScientific) was dried via speed vac and reconstituted in 0.2 M EPPS buffer, pH 8.0. An aliquot of 20 µL of MS grade anhydrous acetonitrile (ACROS Organics) was used to reconstitute 0.5 mg of TMTpro reagent and incubated at room temperature for five minutes with periodic vortexing. The entire 20 µL volume of TMTpro solution was added to the WTC11 or BSA peptide sample. Sample was allowed to incubate at room temperature for 1 hour, with vortexing every 10 min. A 2% aliquot of sample was removed for desalting and was analyzed via LC-MS (TMT labeling quick check method) to check the labeling efficiency. The remaining sample was kept frozen at -80°C until the next day. Samples were treated again with TMTpro if necessary to achieve 90% labeling efficiency. For quenching, sample was thawed, and a 5 µL aliquot of 5% hydroxylamine was added to the sample, vortexed, spun down then incubated for 15 min at room temperature. The sample was then dried via speed vac and reconstituted with 0.1% formic acid in preparation for desalting. After desalting elution, the sample was dried via speed vac and reconstituted with 0.1% formic acid.

#### Mixing of Trigger and Target Peptides

##### BSA benchmarking

A series of TMTpro BSA targets were spiked into 500 ng/µL Jurkat protein digest or HUVEC protein digest with 100 fmol/µL shTMTpro BSA trigger. The BSA target concentrations were 0.1 amol/µL, 1 amol/µL, 10 amol/µL, and 100 amol/µL.

##### AS192 and WTC-11

For eight fraction preparation, a 20 μL (20 pmol) aliquot of shTMTpro-labeled AS192 trigger peptides was added to the tube of quenched TMTpro-labeled WTC11 peptides (100 μg). A 10% aliquot was reserved for Tomahto and DDA analysis of the unfractionated trigger/target mixture. This aliquot was desalted as above and reconstituted in 0.1% formic acid for a final concentration of 1 μg target per μL of sample. The remaining 90% was subjected to high pH reversed-phase fractionation.

#### Offline Fractionation

The 90 μg of TMTpro-labeled WTC11 peptides (endogenous targets) that were spiked with 18 pmol shTMTpro-labeled AS192 triggers were fractionated using the Pierce high pH reversed phase peptide fractionation kit (Thermo Scientific). A total of eight fractions were generated, dried via speedvac, and reconstituted with 0.1% formic acid.

#### LC-MS Methods

##### RP-HPLC Conditions

###### *90* minute RP-HPLC gradient

Desalted sample was analyzed by nanoLC-MS/MS using a Dionex Ultimate 3000 (Thermo Fisher Scientific, Bremen, Germany) coupled to an Orbitrap Eclipse Tribrid mass spectrometer (Thermo Fisher Scientific, Bremen, Germany). An equivalent of 1 μg of peptides was loaded onto an Acclaim PepMap 100 trap column (300 μm x 5 mm x 5 μm C18) and gradient-eluted from an Acclaim PepMap 100 analytical column (75 μm x 25 cm, 3 μm C18) equilibrated in 96% solvent A (0.1% formic acid in water) and 4% solvent B (80% acetonitrile in 0.1% formic acid). The peptides were eluted at 300 nL/min using the following gradient: 4% B from 0-5 min, 4 to 10% B from 5-10 min, 10-35% B from 10-60 min, 35-55% B from 60-70 min, 55-90% B from 70-71 min, and 90% B from 71-73 min, and 4% B from 73-90 min.

###### *4-hour* RP-HPLC gradient

For all samples, injections were loaded onto an Acclaim PepMap 100 trap column (300 µm x 5 mm x 5 µm C18) and gradient-eluted from an Acclaim PepMap 100 analytical column (75 µm x 25 cm, 3 µm C18) equilibrated in 96% solvent A (0.1% formic acid in water) and 4% solvent B (80% acetonitrile in 0.1% formic acid). The peptides were eluted at 300 nL/min using the following gradient: 4% B from 0-5 min, 4 to 28% B from 5-210 min, 28-40% B from 210-240 min, 40-95% B from 240-245 min, and 95% B from 245-250 min.

### Instrument Conditions

#### TMT labeling efficiency check

The Orbitrap Eclipse was operated in positive ion mode with 2.1kV at the spray source, RF lens at 30% and data dependent MS/MS acquisition with XCalibur version 4.5.445.18. MS data acquisition was set up according to the existing method template, “TMT SPS-MS3 RTS.” Positive ion Full MS scans were acquired in the Orbitrap from 400-1600 m/z with 120,000 resolution. Data dependent selection of precursor ions was performed in Cycle Time mode, with 2.5 seconds in between Master Scans, using an intensity threshold of 5e3 on counts and applying dynamic exclusion (n = 1 scans for an exclusion duration of 60 seconds and +/-10 parts per million (ppm) mass tolerance). Monoisotopic peak determination was applied, and charge states 2-8 were included for CID scans (quadrupole isolation mode; rapid scan rate, 0.7 m/z isolation window, 32% collision energy, normalized AGC 100%). MS3 quantification scans were performed when triggered by the real-time search (RTS) algorithm. MS3 (HCD) scans were collected in the Orbitrap with 50,000 resolution, 55% collision energy, Automatic Gain Control (AGC) target of 200%, and custom maximum inject time mode for a maximum inject time of 120 milliseconds (ms) and 10 synchronous precursor selection (SPS) precursors per cycle.

#### DDA settings

The Orbitrap Eclipse was operated in positive ion mode with 2.0 kV at the spray source, RF lens at 30% and data dependent MS/MS acquisition with XCalibur version 4.3.73.11. Positive ion Full MS scans were acquired in the Orbitrap from 375-1500 m/z with 120,000 resolution. Data dependent selection of precursor ions was performed in Cycle Time mode, with three seconds in between Master Scans, using an intensity threshold of 2e4 ion counts and applying dynamic exclusion (n = 1 scans within 30 seconds for an exclusion duration of 60 seconds and +/-10 ppm mass tolerance). Monoisotopic peak determination was applied, and charge states 2-6 were included for HCD MS2 scans (quadrupole isolation mode; 1.6 m/z isolation window, Normalized collision energy at 30%). The resulting fragments were detected in the Orbitrap at 15,000 resolution with Standard AGC target and Dynamic maximum injection time mode.

#### Tomahto API and settings

Tomahto version 1.7.1.29506 was used under an API license agreement with Thermo Fisher Scientific and installed as a software package on the MS instrument computer. The Tomahto software was sourced from the following website: https://gygi.hms.harvard.edu/smartTMTSoftware.html. The target list and analysis parameters were set per user guide instructions using the graphical user interface (GUI). The list of target peptide sequences with corresponding gene annotation was uploaded as a .csv file. The modifications selected for synthetic triggers were the following: shTMTpro (static +313.231 Da) on Lysine (K) and amino terminus of the peptide (NPep), carbamidomethylation (static +57.02146 Da) on Cysteine (C), and oxidation (dynamic +15.9949 Da) on Methionine (M). The modifications selected for endogenous targets were as follows: TMTpro (static +304.2071 Da) on K and NPep, carbamidomethylation (static +57.02146 Da) on C, and oxidation (dynamic +15.9949 Da) on M.

The Orbitrap Eclipse was operated in positive ion mode with 2.0 kV at the spray source and RF lens at 30% using XCalibur version 4.3.73.11. Data acquisition was initiated through Xcalibur, where the instrument method was set to perform only MS1 scans (Orbitrap resolution of 120,000 and mass range 375-1500 m/z) with Standard AGC target and Auto maximum injection time mode. Tomahto, connected through API, ran simultaneously, and was set to monitor MS1 scans for potential trigger m/z within a user-specified mass tolerance (30 ppm), ion intensity threshold (5e4), and charge state match. MS2 scans for trigger and target were acquired in the Orbitrap with an isolation width of 1.0 m/z using HCD or CID (depending on experiment) and 15,000 resolution. Trigger MS2 scans were acquired with 34% collision energy, 120 ms max ion time (IT), and AGC target 1e4. Target MS2 scans were acquired with 34.1% collision energy, 900 ms max IT, and AGC target 1e5. Once real-time trigger/target match criteria were met, as described in Yu et al. 2020, the SPS-MS3 prescan and quantification scans were performed, using default parameters of normal scan mode with AGC target of 1e6 and 10 ms max IT for pre-scan. SPS-MS3 quantification scan parameters were as follows: precursor exclusion window of 5-50 m/z, SPS ion range of 400-2000 m/z, 10 SPS ions selected, SPS ion cutoff of 2% of base peak, MS2 isolation width of 1.2 m/z, MS3 HCD collision energy of 45%, MS3 orbitrap resolution of 50,000, MS3 AGC of 2.5e5, and MS3 max IT of 1000 ms. Default close-out values were used, where a close-out was initiated if three MS3 scans of a trigger are collected and S/N sums to at least 1000. In addition, the “Reset Each Run” checkbox was enabled, which resets the exclusion lists between consecutive runs, so that the full list of target peptides can be tested for each analysis.

### Data Analysis

#### DDA data analysis

Raw files acquired using DDA were searched using Proteome Discoverer (PD) version 2.4.0.305. Raw files from WTC11 or AS192 synthetic peptide experiments were searched against the WTC11 protein database as well as a common contaminants protein database. Raw files from BSA benchmarking experiments were searched against the same contaminants protein database in addition to the Uniprot UP000005640 human proteome with the manual addition of the BSA protein sequence.

The following processing nodes were used: Spectrum Files RC, Spectrum Selector, Sequest HT, and Percolator. Full details for processing and consensus workflow parameters are found in **Supplemental Methods**. In summary, Sequest HT was parameterized as follows: fully tryptic enzymatic digestion with a maximum of two missed cleavage sites allowed, minimum and maximum peptide lengths were set to 6 and 85, respectively, and monoisotopic precursor and fragment ion mass tolerances were set to 15 ppm and 0.05 Da, respectively. In addition, dynamic modifications allowed were the following: TMTpro (+304.207 Da) on peptide N-termini and K, shTMTpro (+313.231 Da) on peptide N-termini and K, Oxidation (+15.995 Da) on M, Acetyl (+42.011 Da), Met-loss (-131.040 Da), and Met-loss+Acetyl (-89.030) on protein N-terminus. Static modification was Carbamidomethyl (+57.021 Da) on C. A concatenated target/decoy strategy was used with validation based on q-value, and a strict Target FDR value of 0.01 was applied. Peptides were further filtered using the Percolator PEP score of ≤ 0.01 using the Peptide Spectral Match (PSM) results file and custom scripts found in the GitHub repository associated with this manuscript.

#### Tomahto data analysis

As a part of the Tomahto method, all synthetic trigger MS2 spectra undergo a real-time peak matching (RTPM) strategy which requires at least six matching experimental fragment ions within +/-10 ppm of theoretical mass.^41^ Once confirmed, trigger MS2 fragment ion and intensities are stored in memory for comparison to corresponding target MS2 spectra. After target MS2 spectra are acquired, they undergo RTPM, and the matching fragment ions are rank ordered according to intensity. SPS fragment ions are selected by meeting the following criteria: (1) b- and y-type ions that have a TMT modification, (2) fragment ion ratios in the target MS2, relative to the highest fragment ion, are within ±50% of the trigger MS2, and (3) at least 50% of the ion signal is attributed to the fragment ion within a 3 m/z window. Quantification scans are performed after these criteria are met.

Additional automatic criteria were met according to the method parameters used. In this study, (1) precursor ion mass tolerance within 10 ppm; (2) fragment ion mass tolerance within 30 ppm; and (3) target retention time matches trigger retention time. For further manual validation, the data analysis module of Tomahto was used to upload the raw files and export the Tomahto results file. The criteria used for further target filtration were the following: (1) ratio of target fragment ions observed over trigger fragment ions observed is at least 50%, (2) allow for cases with 40% fragment ion coverage and 100% isospec purity, (3) isospec purity at least 65%, and (4) relative abundances of top three most abundant SPS fragment ions match between trigger and target MS2. Further analysis was conducted using custom R scripts found in the GitHub repository associated with this manuscript: https://github.com/sheynkman-lab/Alternative-Proteome-Detection-Project.

#### Skyline Data Analysis

Skyline version 23.1.0 was used to generate reports from raw files from Tomahto and DDA experiments. Briefly, transition settings of an ion match tolerance of 30 ppm, minimum m/z of 160, maximum m/z of 3000, method match tolerance of 0.02 m/z, precursor mass analyzer set to Orbitrap, MS1 resolving power of 60,000 at 400 m/z, MS2 acquisition method of PRM, product mass analyzer set to Orbitrap, and resolving power of 15,000 at 400 m/z. Peptide settings were enzyme set to Trypsin [KR|P], static modification of carbamidomethyl on C (+57.021464 Da) with a maximum of five variable modifications and a maximum of one loss. For export of shTMTpro labeled trigger information, structural modifications of shTMTpro K and shTMTpro N-term (both +313.231019 Da) were applied. For export of TMTpro labeled target information, structural modifications of TMTpro K and TMTpro N-term (both +304.207145 Da) were applied. Skyline reports were generated with the following columns: Peptide, Protein, Peptide Modified Sequence, Precursor Mz, Total Area MS1, Peptide Sequence, Best Retention Time, Retention Time, Start Time, End Time, Raw Spectrum Ids, Transition Result Is MS1, Precursor Charge, File Name, Raw Intensities, and Total Area.

#### Peptide-to-protein isoform genome browser track

To map the WTC11 predicted protein isoforms to the UCSC genome browser, the ‘corrected_with_cds.gtf’ file that resulted from running the LRP pipeline mentioned above was run through the GTF2BED module of the GitHub repository associated with this manuscript. Using the other modules in the aforementioned repository, Pogo (https://www.sanger.ac.uk/tool/pogo/) was used to map the predicted AS192 peptides, AS192 peptides detected via DDA, and AS192 peptides detected via Tomahto to genomic coordinates. Final colored BED files were uploaded onto the UCSC genome browser as tracks.

#### Evidence of AS192 in public data repositories

To assess peptide “novelty”, two public MS data repositories, UCSD MassIVE (Mass Spectrometry Interactive Virtual Environment) and PeptideAtlas, were interrogated for evidence of AS192 peptides in previous proteomics experiments. Version 1.3.16 of MassIVE (massive.ucsd.edu) and the Human build of Peptide Atlas (db.systemsbiology.net/sbeams/cgi/PeptideAtlas/Search) were accessed on March 13, 2024. Each of the AS192 target peptide sequences was entered for a manual search of the entire human dataset. Instances where the target peptide sequence was present in either database, in modified or unmodified form, was deemed “annotated”. If a target peptide sequence was absent in either database, in modified or unmodified form, it was deemed “novel”.

To assess the total number of “novel” transcript isoforms identified from WTC11, the results of SQANTI3 isoform classification were used. Isoforms were counted as “novel” if they were annotated as pNIC or pNNC in reference to GENCODE v42.

#### Statistical Software

Data analysis was conducted using R version 4.3.2.^51^ The package ‘tidyverse’ version 2.0.0^52^ was used for data analysis, and visualization through ‘ggplot2,’ and ‘eulerr’ version 7.0.0^53^ was used to create area-proportional Venn diagrams.

## RESULTS AND DISCUSSION

### Defining potential protein isoforms expressed in a human cell line

In this study, we aim to improve analytical detection of alternative protein isoforms through IS-PRM targeting of isoform-specific peptides using the Tomahto software from the Gygi laboratory.^41^

To effectively develop such a method, ideally we need analytical standards of defined mixtures of protein isoforms in a background of complex matrix. However, commercially available proteomics standards, such as Universal Proteomics Standards (UPS) and Proteomics Dynamic Range Standard Set (UPS2), fail to capture the complexity of isoform; therefore, we proceeded to design a model system for the purpose of assay development.

We selected an induced pluripotent cell line, WTC11, reasoning that some prior information about the transcript isoforms expressed in this cell could serve as a first approximation/proxy for protein isoform expression. Such information could be useful to select protein isoforms for targeted MS analysis.

We characterized the transcriptome and predicted isoform-resolved proteome of WTC11 using an LRP pipeline our lab recently developed (**Figure 1A**). Briefly, the transcriptome of the WTC11 cell line was deeply sequenced by collecting long-read RNA-seq data using the PacBio Sequel II platform. The full-length transcriptome was assembled by Iso-Seq 3, annotated for known and novel protein isoforms by SQANTI Protein, and proteins were predicted using the CPAT ORF caller (see **Methods**).

In the WTC11 sample, we predicted 40,846 protein-coding isoforms from 11,755 genes, with an average of 3.5 isoforms per gene. Of the 64K annotated transcripts (GENCODE), 14,559, or approximately 23%, were detected in WTC11, a small fraction of which matched expectation, due to WTC11 representing a single cell population, while the GENCODE transcripts derive from diverse tissues and cell-types.^54,55^ In addition to the annotated transcripts, we found that 24,648 of the 43,828 (56%) detected WTC11 protein-coding isoforms were novel, as assessed via SQANTI Protein (an extension of SQANTI3 developed in the Sheynkman lab).^23,48^

### Selection of target isoform-specific peptides from the proteogenomics database

Using a cell line for long-read RNA sequencing and MS experiments provides insight into the sample-specific transcriptome and predicted proteome, directly informing protein data acquisition. Though the space of candidate isoforms dramatically narrows (e.g., from 300K to a few thousand) with the addition of transcript evidence, there is still an overwhelming number of isoforms which make the scale unfeasible for current MS capabilities. Therefore, in any effort to target previously uncharacterized protein isoforms, the success of target detection is aided by prior weights of the probability of isoforms. Thus, we narrowed our focus to protein isoforms within WTC11 that had the highest likelihood of being expressed and detectable via targeted MS.

First, we examined genes that produced multiple transcript isoforms and, as a consequence, potentially multiple protein isoforms. For all isoforms of a gene, we categorized the most abundant isoform as the major isoform. All other, lower expressed isoforms (at least at the RNA level) are considered minor isoforms. Though the definition of reference and alternative proteins can vary across fields and studies, for our study, we defined any minor isoform as an alternative isoform.

Then, we set out to select alternative isoforms that could reasonably be detected using a bottom-up MS workflow. Therefore, we prioritized cases in which the minor isoform is highly expressed at the RNA level and makes up a non-trivial fraction of the total isoforms expressed for a gene. Frequently, there is one isoform that dominates all expression, and the minor isoform is expressed at negligible, sub stoichiometric levels, which we did not select. Therefore, the abundance of minor isoforms is determined for each gene, whereby isoforms with the highest abundance were preferentially ranked (see **Supplemental Table S1**).

Next, for the candidate alternative protein isoforms, we subjected the predicted protein sequences to *in silico* tryptic digestion to assess the peptides amenability for MS. We compiled major isoform-mapping and minor isoform-mapping peptides between 9 to 20 amino acids in length which were predicted to be retainable on a chromatographic column using AutoRT retention time prediction.^56^ In addition, we ensured the peptide candidates could be chemically synthesized (see **Supplemental Methods**), because a spike-in peptide (trigger) required for Tomahto targeting.

The process for selection of protein isoforms and their associated WTC11 isoform-specific peptides is depicted in **Figure 1A**. The selected peptides, named AS192 peptides, consist of 192 peptides that correspond to 55 genes with one major isoform and at least one minor isoform represented (see **Supplemental Table S1**).

### Implementation and benchmarking of Tomahto IS-PRM

Before analysis of the isoform-specific peptides, we first evaluated the performance of Tomahto of a well-defined mixture of BSA protein digest. First, we installed Tomahto using the Thermo API on our Eclipse (see **Methods**). The assay configuration of IS-PRM, and specifically Tomahto, requires labeled trigger peptides of interest in combination with differently labeled target peptides. The trigger peptides were labeled with super heavy TMTpro (shTMTPro), and target peptides were labeled with the standard TMTpro. As Tomahto is a type of IS-PRM, MS1 ions are monitored for the presence of user-defined trigger peptide masses (**Supplemental Figure S1**). Considering that the trigger and corresponding target peptides are from the same amino acid sequence and their chemical labels are isobaric, they will undergo the same chromatographic separation. However, the different labeling strategies between trigger and target, instrumental in TOMAHAQ and Tomahto, result in distinct gas-phase separation. When a precursor trigger m/z is detected within ± 10 ppm at ≥ 50,000 ion counts and six trigger-derived fragment ions within ± 30 ppm are matched to the predicted trigger spectra, the Tomahto interface triggers MS2 acquisition of the corresponding target m/z at the appropriate offset m/z (9 Da/charge state of the peptide).

Many factors could influence analysis of low abundance isoform-specific peptides in complex mixtures. Before analyzing AS192, we first determined the sampling sensitivity exhibited by Tomahto versus compared to standard DDA mode. Specifically, we compared the peptide detection rates when using DDA versus Tomahto mode, testing known concentrations of trigger and target peptides using isobarically labeled BSA peptides. We used a high, constant concentration of the triggerant, and vanishingly low amounts of target peptide. Accordingly, we analyzed samples in triplicate in a dilution series (0.1 to 100 amol on-column) of BSA target peptides combined with 100 fmol/µL of trigger BSA, within a complex matrix of cell lysate peptide digest (**Figure 2A**).

DDA runs were searched against a FASTA combining a human UniProt proteome (UP0000056400) database with the addition of BSA. Peptides passing a 1% FDR and a Percolator PEP score of ≤ 0.01 were considered identified. Although over 24,635 distinct peptides were identified across the three replicate runs, we identified just one BSA tryptic peptide at or below 100 amol (**Figure 2B**).

The same injected samples run in Tomahto mode returned identifications for 12 of the 27 BSA peptides at these same low concentrations. Notably, we were able to detect BSA peptides using Tomahto at 0.1 amol across multiple runs, demonstrating the sensitivity of this method. For both these DDA and Tomahto runs, MS data was collected using HCD fragmentation. Based on the promising results of Tomahto, we tested CID fragmentation within the Tomahto method because of its use as the preferred fragmentation method in the original Tomahto manuscript.^41^ We observed a marked increase in target coverage between the use of HCD fragmentation (12/27 BSA peptides) and CID fragmentation (20/27 BSA peptides) (**Figure 2C**).

**Figure 2.**
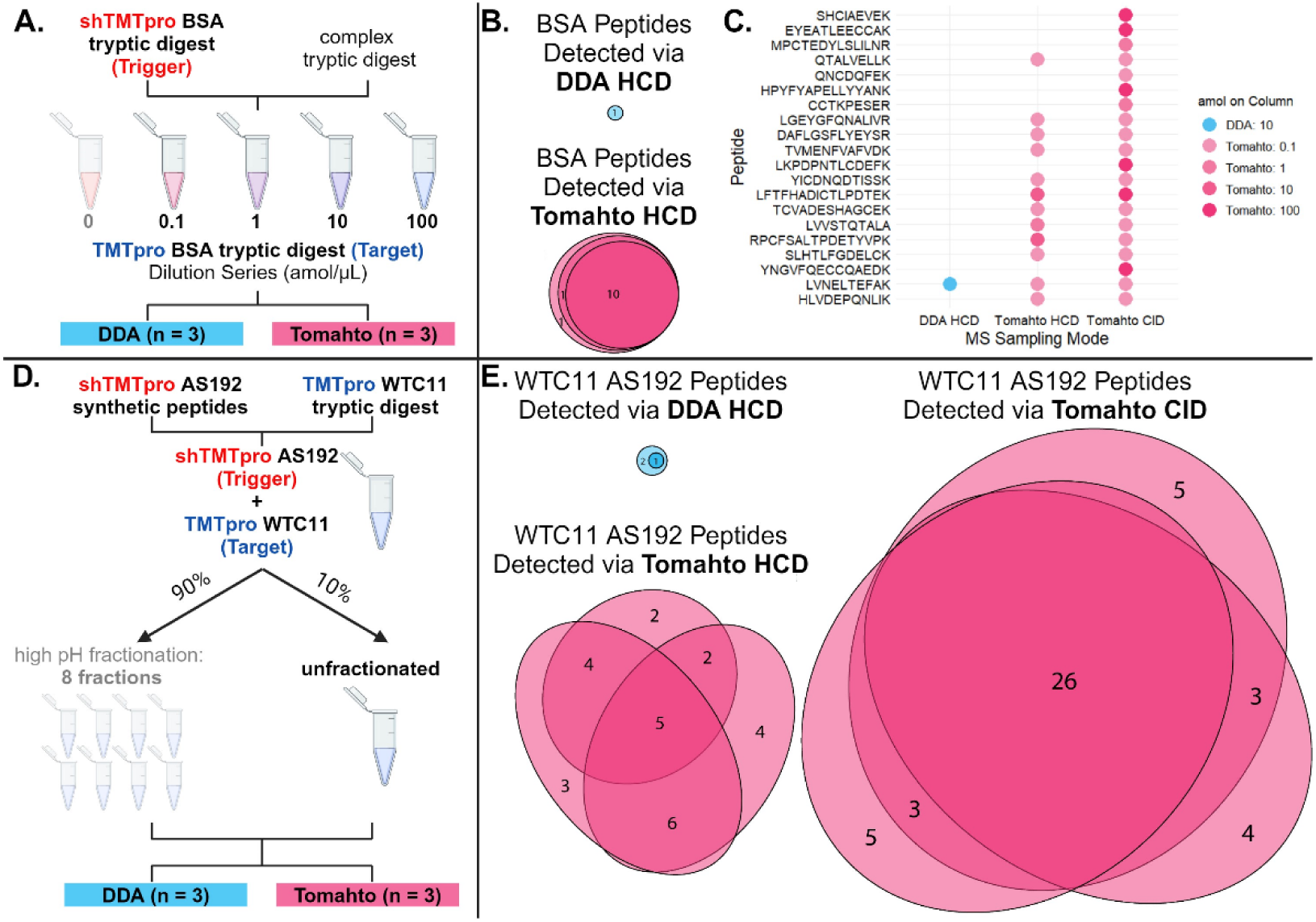
Assessment of Tomahto repeatability and peptide detection rates. A) Schematic of BSA benchmarking experiment. B) Detection of TMTpro-labeled BSA peptides in complex matrix over three replicates via DDA with HCD (top) and Tomahto with HCD fragmentation (bottom). C) Detected BSA peptides from (B) with additional peptides detected via an analysis of the same sample using Tomahto with CID fragmentation. Shading represents the lowest amol of TMTpro BSA on column at which the peptide was detected. D) Schematic of experiment for detecting endogenous isoform-specific peptides (AS192) in WTC11 using Tomahto or DDA. E) Number of AS192 peptides detected in unfractionated WTC11 across triplicate injections using DDA HCD (top left), Tomahto HCD (bottom left), or Tomahto CID (right). BSA = bovine serum albumin, DDA = data dependent acquisition, HCD = higher energy collisional dissociation, CID = collision-induced dissociation.

### Tomahto targeted detection of isoform-specific peptides

After establishing Tomahto within our lab as a tool for detecting previously observed target peptides, we proceeded to characterize alternative proteins in WTC11, by using the AS192 peptides. As a first step in assessing the feasibility of detecting endogenous AS192 sequences using Tomahto, we performed DDA and Tomahto experiments on neat injections of 1 μg of pooled shTMTpro labeled AS192 synthetic peptide triggers **(Supplemental Table S2).** The goal of this step was two-fold: first, to determine which AS192 synthetic peptides were detectable via DDA, and second, to elucidate any instances of contamination with TMTpro reagents that could confound results. Conducting DDA on the neat AS192 triggers resulted in the detection of 104 shTMTpro labeled AS192 trigger peptides and no detection of TMTpro labeled targets, as anticipated. Conducting the experiment using Tomahto resulted the detection of 158 triggers and the aberrant detection of five “target” AS192 peptides: NSGQGCIGG, VPAQPAAEQR, HGGCLLQESR, SPSQGSPIQSSD, and RPASLGCGGWLLPGR. However, inspection of the quantification of the TMTpro tag within the Tomahto interface yielded no signal in the expected channels. Therefore, the detection of these “targets” via Tomahto were determined to be false positives. In subsequent analyses in WTC11 protein digests in which the endogenous (target) peptides were detected, manual inspection of the quantification channels was performed to ensure that the targets did indeed contain TMTpro labels. In total, 174 out of the 192 shTMTpro triggers were detected upon DDA and Tomahto assessment, with 16 trigger peptides detected exclusively in DDA, and 12 trigger peptides detected only via Tomahto.

Next, Tomahto was applied towards the detection of isoform-specific peptides in the WTC11 sample (endogenous AS192 peptides). **Figure 1B** depicts the experimental workflow. Briefly, shTMTpro-labeled synthetic trigger AS192 peptides were spiked into TMTpro-labeled WTC11 peptide digest. The sample was then either left unfractionated or underwent high-pH separation into eight fractions and injected in triplicate (**Figure 2D**). Analysis of the unfractionated WTC11 sample using DDA (see **Methods**) resulted in identification of 12,386 peptides and 3,415 protein groups. Analysis of the fractionated samples resulted in 64,778 peptides and 8,673 protein groups.

In the unfractionated WTC11 sample, a total of 26 peptides were identified using Tomahto HCD, with 10 peptides detected in all three replicates (**Figure 2E**). In contrast, Tomahto using CID fragmentation (Tomahto CID) identified a total of 46 peptides, with 26 peptides detected in all three Tomahto CID replicates. This is a significant improvement over DDA, which only captured three total peptides, none replicating across all three injections.

While it was expected that DDA would detect fewer target peptides, it was initially not expected for CID fragmentation mode to detect more peptides than HCD mode. In contrast to our findings, a previous study using DDA comparing HCD and CID using Orbitrap detection concluded that HCD, not CID, resulted in more protein identifications.^57^ This might be due to CID producing fewer but more intense fragments with corresponding larger m/z values, which make these fragments more amenable to SPS selection and MS3 quantitation via the Tomahto interface^41^. In contrast, HCD tends to result in more fragmentation events with subsequently lower m/z and intensity values.^58^ These results indicating that using Tomahto CID results in a roughly 2-fold increase in peptide identifications over Tomahto HCD, combined with the previous literature on HCD outperforming CID in DDA experiments, led us to conduct our fractionation experiments using DDA HCD and Tomahto CID.

Interrogation of the fractionated WTC11 samples using Tomahto CID resulted in the identification of 65 out of the 192 AS192 target peptides. In contrast, 21 DDA HCD peptides were identified, yielding a greater than 3-fold increase in peptide identifications when using Tomahto CID. In both unfractionated and fractionated WTC11 peptide sample and across different fragmentation modes, Tomahto returned a higher number of confirmed peptide identifications (**Figure 3A**).

### Tomahto provides evidence of major and minor protein isoforms

Having established the use of Tomahto to increase rates of identification of endogenous AS192 peptides in WTC11 compared to DDA, we proceeded to characterize the protein isoforms supported by the i peptides.

**Figure 3.**
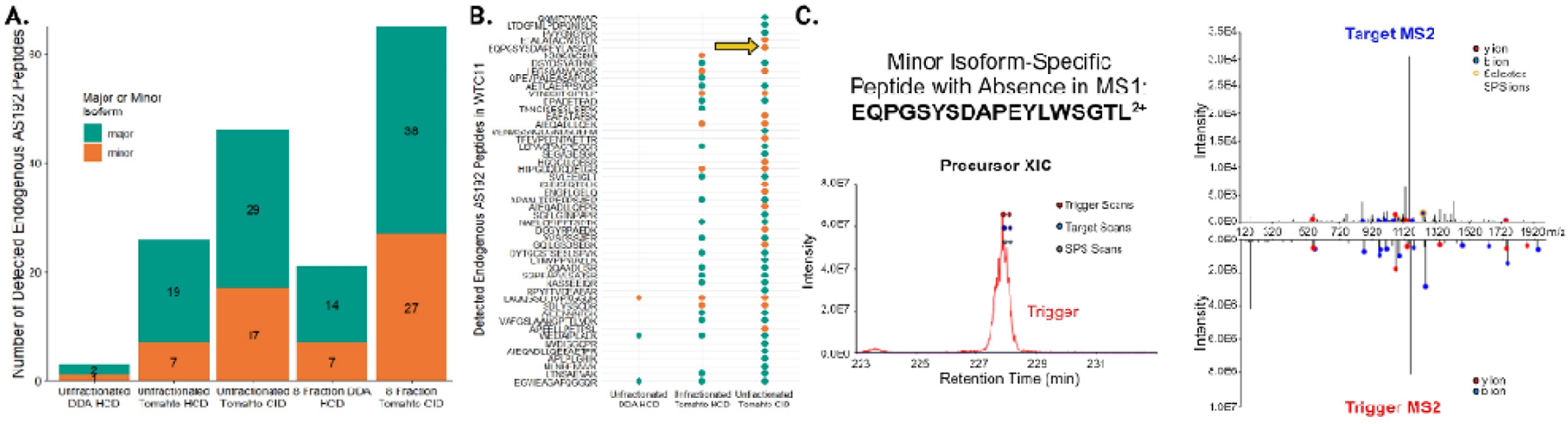
Tomahto detects more isoform-specific peptides than DDA, even when the precursor is not detected in MS1. A) Total number of AS192 peptides detected per experiment, separated by whether they inform on major or minor isoform presence (see **Table 1**). B) Detection of endogenous isoform-specific peptides (AS192) in unfractionated WTC11 by DDA and Tomahto with HCD fragmentation and CID fragmentation. Color indicates whether the peptide is specific to a major isoform (teal) or minor isoforms (orange). Arrow indicates a peptide precursor that was not detected in MS1 but was captured in MS2 using Tomahto. C) Extracted ion chromatogram depicting minor isoform peptide precursor EQPGSYSDAPEYLWSGTL^2+^ and the associated mirror plot of the fragmentation for the trigger peptide (bottom) and corresponding endogenous peptide (top). Images obtained and adapted from the Tomahto software interface.^41^

Individual peptide sequences and their detection across unfractionated WTC11 experiments are shown in **Figure 3B**. Use of Tomahto in the interrogation of AS192 endogenous peptides in unfractionated WTC11 yielded an appreciable increase in identifications from one minor isoform-specific peptide found via DDA, to seven using Tomahto with HCD fragmentation, to 17 identifications using Tomahto with CID fragmentation. By definition, minor isoforms are lower in transcript abundance compared to their major counterparts and are theoretically more analytically challenging to identify.^59^

Notably, we detected an endogenous minor isoform-specific peptide lacking MS1 precursor signal: EQPGSYSDAPEYLWSGTL (see arrow in **Figure 3B**). Corresponding MS1 precursor chromatogram and MS2 fragmentation for both trigger and target are shown in **Figure 3C**. The fact that MS2 acquisition scans are initiated by detection of a trigger peptide, not precursor MS1 signal, increases sensitivity to detect low abundance peptides, which can be critical for detection of lower abundance minor isoforms.

However, the use of trigger-based targeting is a double-edged sword. While use of triggers allows for MS2 detection of targets without corresponding MS1 detection, it requires sufficient MS1 precursor intensity and MS2 fragmentation of the trigger for detection of the target. Selection of peptides for trigger-based targeting is still limited to the selection of peptides that fly, ionize, and fragment sufficiently. For example, peptide ENLLVEDSLMIECSAR was detected in two out of three replicate runs of fractionated WTC11 using DDA but was not detected in any Tomahto runs (**Table 1**). Upon investigation of the Tomahto output files in the relevant fraction, it was found that the MS2 trigger was acquired, but the trigger spectra did not “Pass” the internal Tomahto criteria for subsequent triggering of target acquisition (6 fragment matches ± 10 ppm of theoretical, ≥ 50,000 ion counts) (**Supplemental Figure S2**). Therefore, even though the endogenous ENLLVEDSLMIECSAR peptide was likely present in the sample, Tomahto failed to detect it.

While Tomahto returned significantly more peptide identifications than DDA, it did not return identifications for 115 out of the 192 peptides that were included in these experiments. As stated previously, upon investigation of the triggers by themselves via DDA and Tomahto, we detected a total of 174 peptides. All the peptides that were identified over the course of this study had a corresponding detectable trigger. However, that means that the remaining 96 AS192 target peptides remain undetected. One potential reason is that it is conceivable that the most abundant form of the protein isoform within our model system contains one or more post-translational modifications (PTM). Inherent to IS-PRM methods is that the trigger m/z must correspond to the expected target m/z for subsequent identification to occur. While we did incorporate dynamic methionine oxidation within the Tomahto interface for both trigger and target, other possible PTMs on targets were not included. Another potential reason for this is that the detected transcripts might not have been translated into stable proteins.^60^ While there is a correlation between transcript number and protein expression,^61^ this relationship is not one-to-one. Until the protein itself is detected, it cannot be assumed based on transcript information alone that it exists, which highlights the importance of conducting paired isoform-resolved transcriptome and proteome studies. Another potential reason for the lack of identification is the overall reduced proteotypicity of the peptides that were not detected. Proteotypicity^62^ is defined as the likelihood of a peptide to be detected via MS and is reliant on multiple factors including ionization, hydrophobicity, and susceptibility to enzymatic digestion. Using the Prosit proteotypicity score (Mathias Wilhelm, unpublished), we assessed the proteotypicity of the AS192 peptides. While there was no difference in proteotypicity scores (t-test; p = 0.9113) between peptides originating from major (3.82 ± 4.61 SD, n = 83) and minor (3.90 ± 4.65 SD, n = 110) protein isoforms, there was a statistically significant difference (t-test; p = 0.0119) in the mean proteotypicity score between peptides detected (4.87 ± 4.48 SD, n = 78) versus not detected (3.18 ± 4.61 SD, n = 115) in this study (**Supplemental Figure S3**).

**Table 1.**
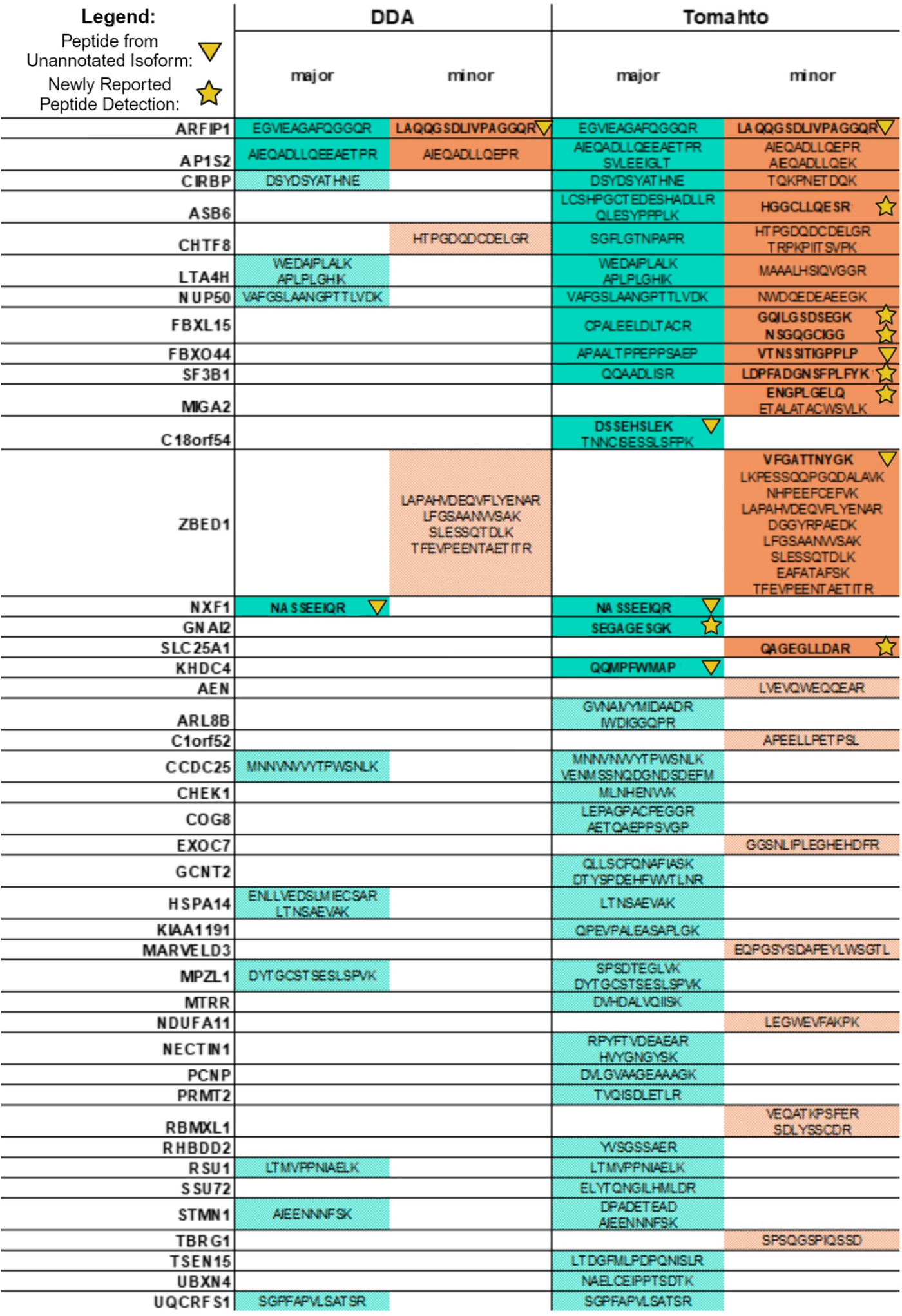
Major and minor isoform-specific peptides detected in unfractionated and fractionated WTC11 using DDA or Tomahto (n = 3 injections). Peptides are grouped by the gene they are associated with. Peptide sequences that are labeled with an upside-down triangle derive from isoforms that were not annotated in GENCODE v42. Peptide sequences that are labeled with a star indicate peptides that had not been reported as detected in mass spectrometry repositories.

**Table 2.** Summary of findings comparing the detection of endogenous AS192 peptides in WTC11 via DDA or Tomahto (see also **Supplemental Table S2**).

|  | All<br>Predicted<br>AS192 | Total<br>Detected<br>AS192 | Detected<br>AS192 via<br>DDA | Detected<br>AS192 via<br>Tomahito |
| --- | --- | --- | --- | --- |
| Peptides | 192 | 78 | 21 | 77 |
| Major Isoform-Specific Peptides | 82 | 46 | 14 | 45 |
| Minor Isoform-Specific Peptides | 110 | 32 | 7 | 32 |
| Isoforms | 114 | 54 | 16 | 54 |
| Genes | 55 | 43 | 14 | 43 |
| ★ Newly Reported Peptide Detection | NA | 8 | 0 | 8 |
| ▼ Peptide from Unannotated Isoform | 21 | 5 | 1 | 5 |

### Tomahto informed by long-read sequencing provides insight into protein isoform expression

To annotate the alternative isoforms supported by detected peptides, we created a custom UCSC Genome Browser track^63^ to visualize RNA transcripts, their predicted proteins, and the associated isoform-specific peptides. Such visualizations provide genomic and alternative splicing context of the peptide identifications (**Figure 4**).

Of note, we captured pairs of major and minor isoform-specific peptides corresponding to the genes *AP1S2*, *ARFIP1*, *CIRBP, ASB6, CHTF8, LTA4H, NUP50, FBXL15*, *SF3B1*, and *FBXO44* (**Table 1**). The identifications of all sets of major and minor isoform pairs except for *ARFIP1* and *AP1S2* were exclusive to Tomahto. Detection of peptides from *FBXO44* and *SF3B1* are shown in **Figure 4A** and **4B**. Of particular interest is the alternative isoform found from the gene *SF3B1*. This gene encodes the largest subunit of the core complex in the spliceosome, which mediates alternative splicing.^64^ Mutations of *SF3B1* occurring in malignancies are correlated with adverse patient outcomes. It has been posited that *SF3B1* mutated proteins might serve as a biomarker in patients,^64^ making detection of specific isoforms of *SF3B1* an important avenue of research. By using methods like Tomahto, we can confirm stably expressed isoforms arising from splicing variations in pathological or physiological states.

Additionally, we detected five peptides from previously unannotated isoforms, defined as being found in the WTC11 proteome but not annotated in the GENCODE v42 reference proteome (**Table 1**). *C1orf52* (**Figure 4C**) is an understudied protein coding gene, whose variants have been linked to risk of multiple sclerosis.^65^ *C1orf52* has been identified as an RNA binding protein,^66^ which are known to be pivotal in regulating gene expression, with mutations in these proteins linked to a host of pathologies.^67^

To assess MS detectability of our AS192 target peptides, we cross-checked the UCSD MassIVE (https://massive.ucsd.edu) and PeptideAtlas^68^ MS data repositories for evidence of AS192 peptide observation in previous proteomics experiments. A total of 141 of the AS192 peptides had evidence of previous MS observation, while 51 were absent from these databases (**Table 2**). Using Tomahto, we detected eight previously undetected peptides including the minor isoform-specific peptide derived from *SF3B1* (**Table 1**). Detection of previously unannotated isoforms and peptides which had not been reported in mass spectrometry data repositories highlights the power of coupling long-read sequencing-informed predicted protein databases with targeted MS. Although *de novo* peptide sequencing via MS has made significant advances,^69^ the vast majority of MS experiments require *a priori* knowledge of potential protein sequences. By informing our targeted approaches with sample-specific proteome prediction and using synthesized peptide internal standards to bolstering confidence in spectral matches, our approach should increase the rate of and confidence in discovery of alternative protein isoforms.

**Figure 4.**
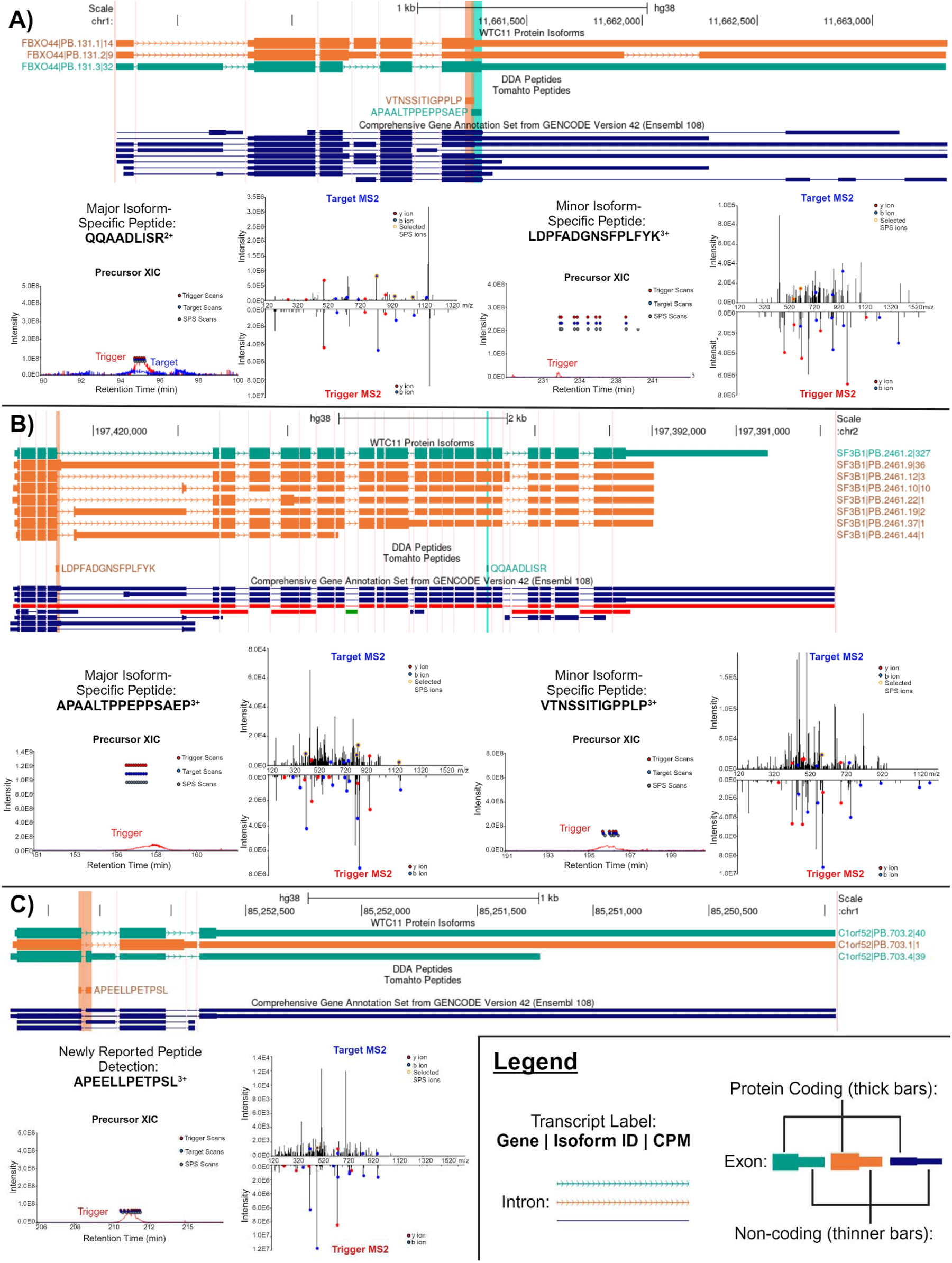
Examples of MS evidence for co-expression of major and minor isoforms as well as detection of unannotatedisoforms. Each example contains a figure panel of a UCSC Genome Browser image with three tracks. The top track displays the long-read RNA-seq-predicted protein isoforms, differentiated as the major (teal) or minor (orange) isoforms by color. The middle track displays the isoform-specific peptides for the major (teal) and minor (orange) isoforms. The bottom track displays GENCODE v42 annotated proteins. Underneath the Browser image are the extracted precursor ionhromatograms and corresponding MS2 results for the trigger peptide (bottom) and endogenous target peptide (top). A) Predicted *FBXO44* protein isoforms and corresponding major isoform-specific peptide APAALTPPEPPSAEP^3+^ and minor peptide VTNSSITIGPPLP^3+^. B) Depiction of *SF3BP1* transcript isoforms and corresponding major isoform-specific peptide QQAADLISR^2+^ and minor peptide LDPFADGNSFPLFYK^3+^. C) Depiction of C1orf52 minor peptide APEELLPETPSL^3+^ which maps to an unannotated isoform.

## CONCLUSIONS

Combining sample-specific long-read RNA sequencing and Tomahto targeted MS is a powerful tool to assess the AS protein isoform landscape. This method allows for the “targeted discovery” of isoforms that were detected via transcriptomics whose protein product was yet unobserved. As a proof of concept, Tomahto outperformed DDA in the overall detection of isoform-specific peptides, identification of multiple protein isoforms derived from a single gene, and detection of unannotated isoforms and peptides which had not been reported in mass spectrometry data repositories. These new observations support the hypothesis that overcoming technical challenges can help improve MS coverage of the alternative proteome. By confirming protein expression of alternative isoforms, this provides an avenue for future studies into their biological roles, with the potential for identification of novel biomarkers or prognostic indicators.

While using an IS-PRM method provides a robust way of detecting target peptides, it does come with limitations. Namely, the generation of synthetic trigger peptides can be cost-prohibitive for some labs. Additionally, a false discovery rate is difficult or impossible to calculate for these experiments. While we applied rigorous filtering criteria when assigning peptide identifications, we acknowledge that false positive identifications can still occur. Therefore, future studies implementing this method of “targeted discovery” for the purposes of validating candidate proteins should likely conduct orthogonal methods using a non-trigger-based approach. Additionally, while the use of synthetic triggers affords multiple benefits, including empirical evidence of the fragmentation pattern and retention time of a peptide of interest in the unique biological system of study, there is a small, but not insignificant, chance of the trigger peptide being “identified” as your target. While the 9 m/z offset trigger mitigates this risk, scrutiny of the reporter ion quantification channels to ensure the expected tag is captured is vital to increase confidence in identifications.

In the future, comparisons of Tomahto using PRM with retention time scheduling, DIA-based SWATH MS,^70^ and the trigger-free advanced targeting method GoDig^38^ will serve to add more context to how Tomahto compares to standard targeting methods and newer MS techniques, respectively. By applying the combination of long-read proteogenomics and internal standard parallel reaction monitoring (LRP-IS-PRM), we can begin to systematically explore the alternative proteome under different conditions. For example, isoform switching, whereby a gene expresses one isoform under certain conditions and a different isoform under separate conditions, is generally investigated at the RNA transcript level and on a case-by-case basis on the protein level. This phenomenon is found in a wide array of physiological and pathological states, ranging from cellular differentiation^71^ to cancer.^72^ By leveraging LRP-IS-PRM, these differential splicing events can be quantified at the protein isoform level on a larger scale, with the inclusion of multiplexed experimental groups for direct comparisons. This study is the first to successfully combine proteogenomic approaches with the trigger-based peptide targeting software Tomahto to interrogate the alternative proteome.

## SUPPORTING INFORMATION

R and Python Scripts used to process or analyze WTC11 and GENCODE isoforms and predicted tryptic peptides, Proteome Discoverer and Tomahto results files, and generating UCSC browser tracks are found in the following Github repository: github.com/sheynkman-lab/Alternative-Proteome-Detection-Project. Raw sequencing data files can be found in the National Institutes of Health (NIH) National Library of Medicine’s Sequence Read Archive with the project accession number PRJNA1090880. Mass spectrometry raw files, Proteome Discoverer PSM result files, and Tomahto result files have been deposited to the ProteomeXchange Consortium via the PRIDE^73^ partner repository with dataset identifiers PXD050904 and PXD050909. Additional data files generated from this study are available upon reasonable request to the corresponding author.

## ACKNOWLEDGEMENTS

We thank Leon Sheynkman for proteomics sample preparation, Madison Mehlferber for long-read proteomics pipeline work, and Jerryd Meade for RNA sample preparation. We also thank Dr. Qing Yu for consultation regarding Tomahto installation and application for this study. This study was supported by the National Institute of General Medical Sciences at the National Institutes of Health (NIH) under R35GM142647 to GMS. Support to JAK provided by NIH T32 HL007284 Cardiovascular Research Training Program. Figures created using Biorender.

## AUTHORS’ CONTRIBUTIONS

GMS: conceived of the project, designed the study, supervised personnel JAK, EDJ, SB, BTJ, ML, and EFW, manuscript writing and review. JAK: experimental design, data collection, data analysis scripts, data validation, manuscript lead. EDJ: experimental design, data collection, data validation, manuscript writing and review. SB: Long-read proteogenomics and isoform analysis. BTL: Long-read proteogenomics analysis. EFW: Long-read proteogenomics analysis, Genome Browser tracks, data visualization. ML: data collection and manuscript review. AF: WTC11 cell culture. MW: procurement of Prosit proteotypicity scores for AS192 peptides.

