## Supplemental Figures for "IS-PRM-based peptide targeting informed by long-read sequencing for alternative proteome detection"

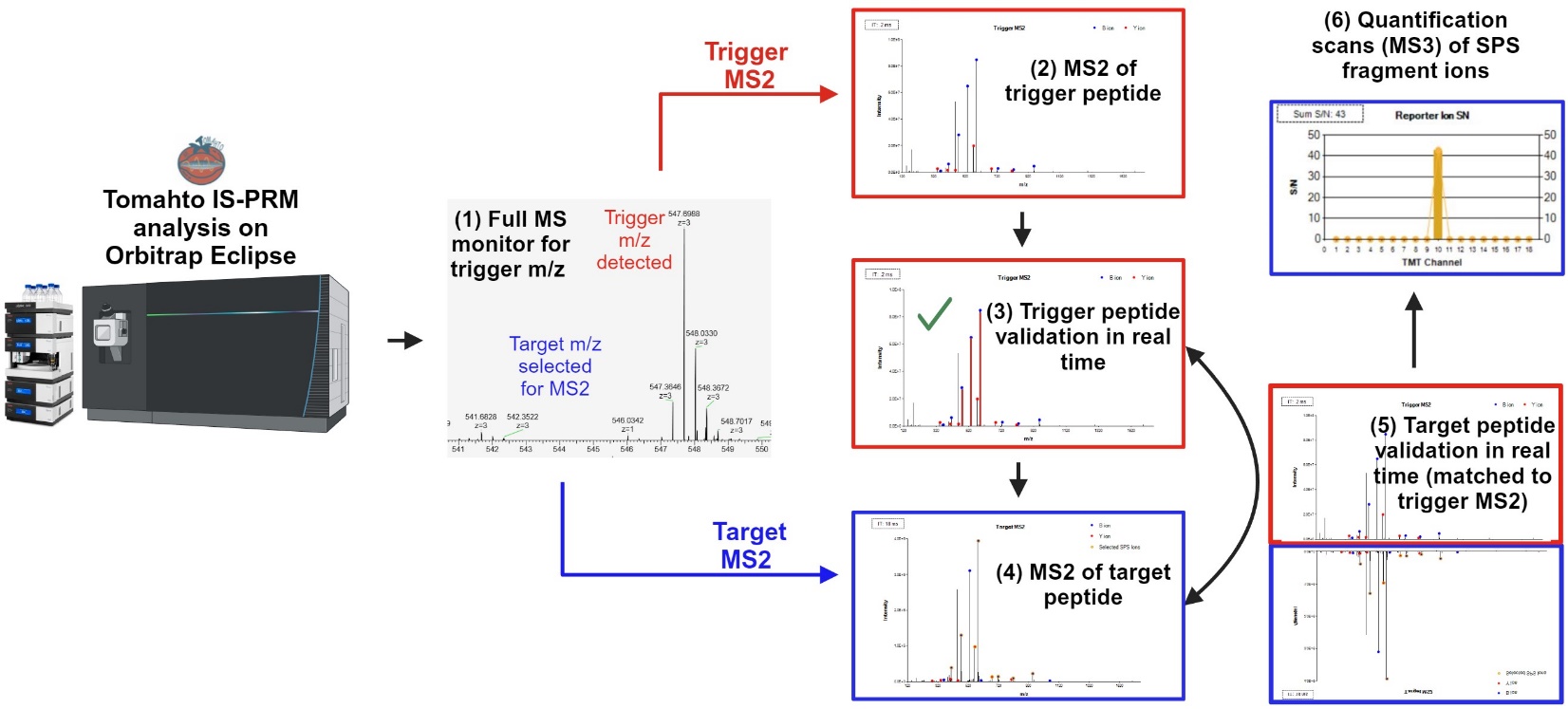

**Supplemental Figure S1.** Tomahto API-controlled instrument method overview. (1) Tomahto operates in "observation" mode to monitor for the presence of shTMTpro-labeled synthetic trigger peptides (user-input via Tomahto GUI). MS1 scans are acquired in the Orbitrap with 60,000 resolution. (2) Upon trigger detection, a trigger MS2 scan is performed and detected in the Orbitrap. (3) As a part of the Tomahto method, all synthetic trigger MS2 spectra undergo a real-time peak matching (RTPM) strategy, as described in Yu et al. 2020, which requires at least 5 matching experimental fragment ions within +/- 30 ppm of theoretical mass (user defined). (4) Once confirmed, an MS2 scan of the target peptide m/z is acquired. Trigger MS2 fragment ion and intensities are stored in memory for comparison to corresponding target MS2 spectra. (5) After target MS2 spectra are acquired, they undergo RTPM and the matching fragment ions are rank ordered according to intensity. SPS fragment ions are selected by meeting the following criteria: b- and y- type ions that have a TMT modification; fragment ion ratios in the target MS2, relative to the highest fragment ion, are within ±50% of the trigger MS2; at least 50% of the ion signal is attributed to the fragment ion within a 3 m/z window. (6) Quantification scans (SPS-MS3) are performed after these criteria are met.

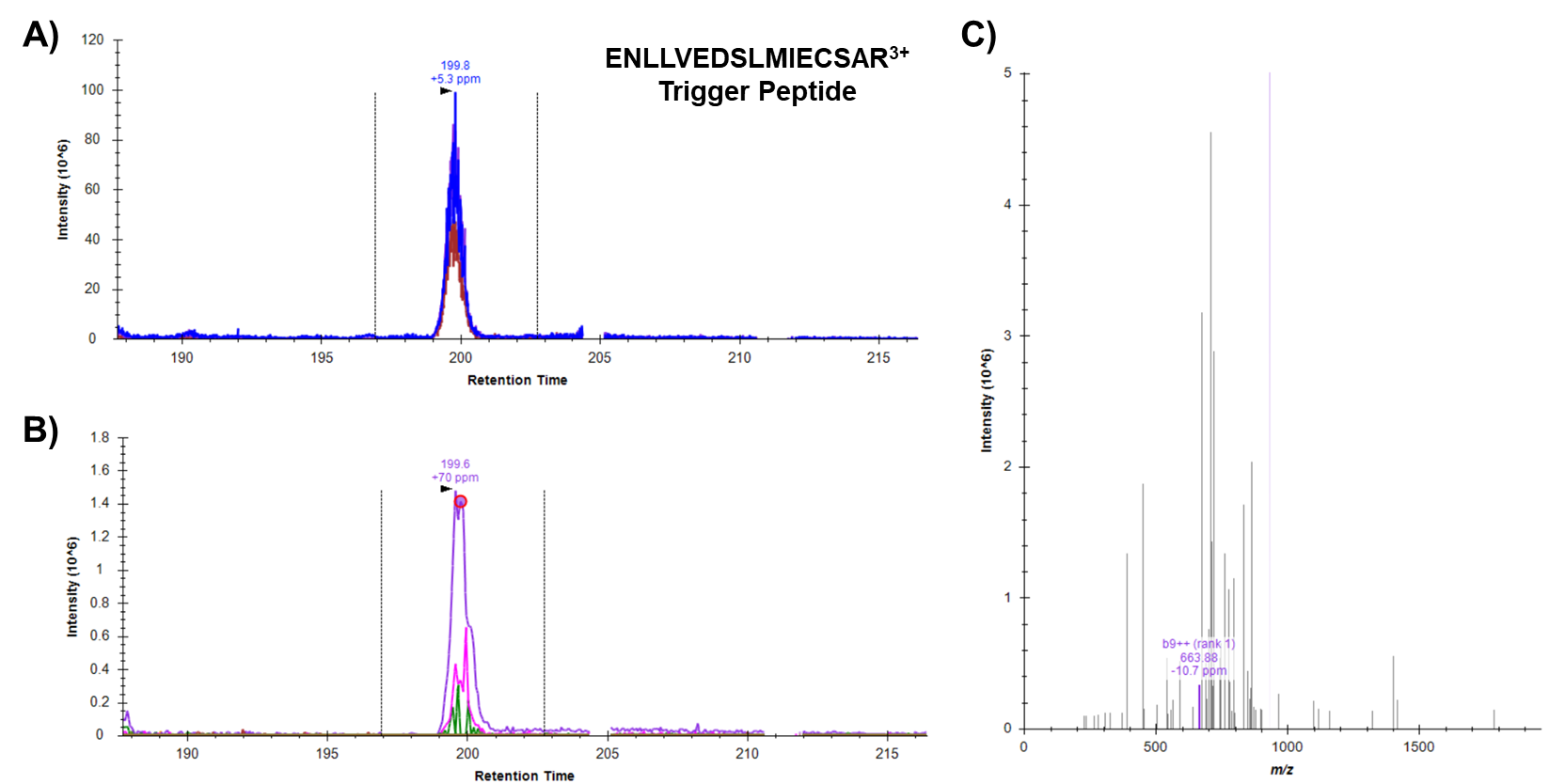

**Supplemental Figure S2.** Investigation of ENLLVEDSLMIECSAR^3+^ trigger peptide via Skyline. A) Extracted ion chromatogram depicting the presence of trigger peptide precursor ENLLVEDSLMIECSAR^3+^ from 8 Fraction Tomahto CID replicate 3. B) Corresponding MS2 acquisition of the trigger peptide. C) Fragmentation of the trigger peptide.

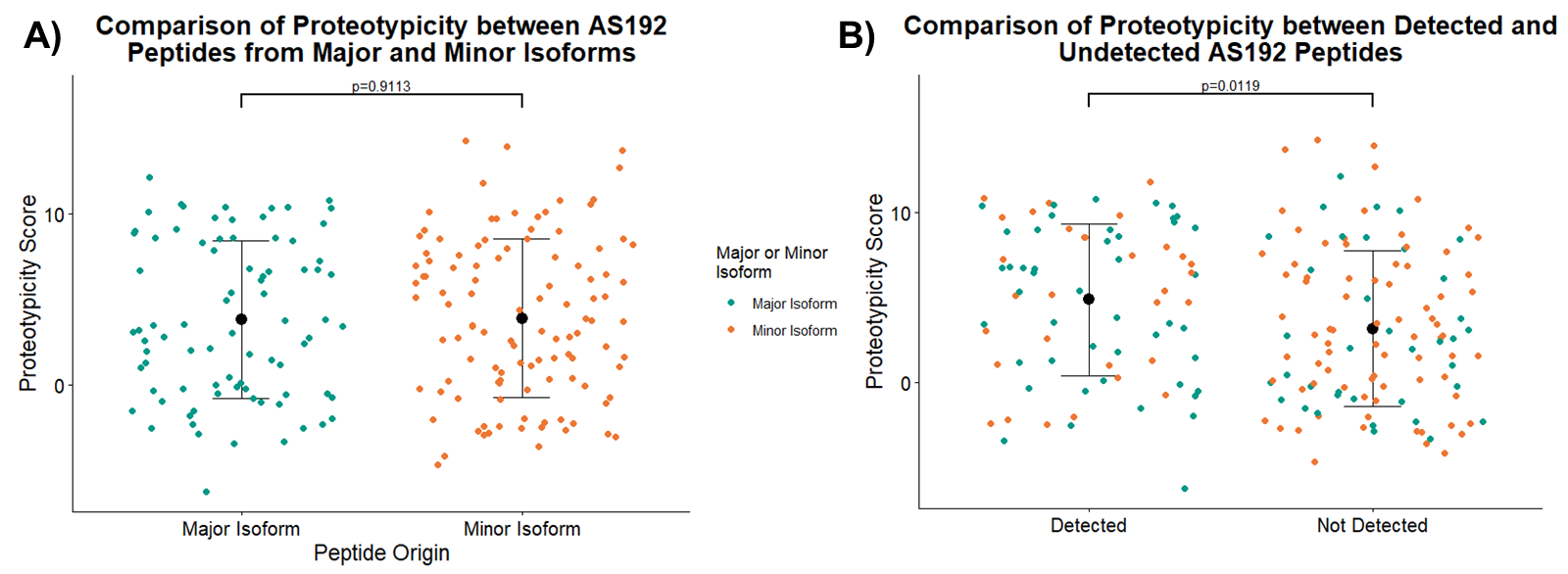

**Supplemental Figure S3.** Assessment of the Prosit predicted proteotypicity scores between AS192 peptides. Mean and standard deviation of proteotypicity score by group depicted by the larger black point and error bars, respectively. The p values were calculated between groups by conducting Welch’s t tests with unpooled variance, α = 0.05. Overlayed points depict the individual proteotypicity scores for each AS192 peptide, colored by major or minor isoform origin. A) Comparison of proteotypicity score of AS192 peptides grouped by whether the peptides originated from a major isoform (n = 83 peptides) or a minor isoform (n = 110 peptides). B) Comparison of proteotypicity score of AS192 peptides grouped by whether the peptides were detected (n = 78 peptides) or not detected (n = 115 peptides) over the course of this study.
