## Supplemental Methods for "IS-PRM-based peptide targeting informed by long-read sequencing for alternative proteome detection"

***Details of AS192 peptide selection***

Selection of target peptides

The LRP pipeline was applied to the WTC11 sample using long-read RNA sequence information. The output protein database was subjected to in-silico tryptic digestion (no missed cleavages, 9-20 amino acids in length). The resulting peptides were filtered to retain ones that were isoform-specific and whose major gene isoform had a transcript abundance of at least 10 full length counts per million mapped (CPM). CPM is used as a normalized measure of transcript abundance, such that for every 1,000,000 RNA transcripts present in a given cell, a defined number of copies came from a particular transcript.[^1^](https://sciwheel.com/work/citation?ids=11601692&pre=&suf=&sa=0&dbf=0) In our group we have observed that, at the gene level, transcripts above 10-25 CPM are more likely to be detected via MS.[^2^](https://sciwheel.com/work/citation?ids=12599088&pre=&suf=&sa=0&dbf=0)

Our peptides were categorized as being specific for either the major gene isoform or an alternate isoform (minor gene isoform). Further filtering was done to retain the peptides from genes that contained both major isoform **and** at least one minor isoform-specific peptide.
Peptides from 79 genes passed these filtering steps, with 334 major isoform peptides (105 of which had been identified from DDA analysis) and 140 minor isoform peptides (22 of which had been identified from DDA analysis). These 474 isoform-specific peptides were the targets for MQL analysis. The majority of the WTC11 target peptide list had not been previously observed in a DDA run.

Of the 474 peptides, we wanted 192 for synthesis (to fit 2 96-well plates). To filter further, minor gene isoform peptides were sorted by abundance. Then the gene with the most abundant minor isoform was selected. All minor isoform peptides mapping to that gene were selected. In addition, 2 major isoform peptides were included (where possible). This was repeated until at least 192 peptides were selected. The final list was derived from 55 genes and contained 82 major isoform-specific peptides and 110 minor isoform-specific peptides.

Normalization of synthetic trigger peptide abundance

Initial normalization of synthetic trigger peptides was conducted by pooling 10 µL of each peptide together into a final 1920 µL solution. The pool then underwent serial dilutions of 1/10, 1/100, and 1/1000 the original pool concentration. Dilutions were completed with 0.1% formic acid (FA). Peptides were analyzed with the Orbitrap Eclipse Tribrid instrument and the peak height of each peptide was recorded using FreeStyle.

A second round of normalization was conducted by selecting the dilution factor that placed each peptide closest to a peak height between 1e7 and 1e8. Peptides that fell below this range at a 1/10 dilution were assigned a dilution factor of 1. Peptides were pooled together by their assigned dilution factor (10 µL each) and diluted accordingly. All 192 peptides were pooled into one sample; the volume contribution of each dilution group was based on the percentage of the total peptides that belonged to each group. MS injection and FreeStyle data analysis was completed as before.

The final AS192-norm sample was prepared by selecting an individualized dilution factor for each trigger peptide. Peptides with a peak height between 1e7 and 1e8 did not change. New dilution factors were selected based on estimations from previous data to bring peak height within range. Dilutions of each peptide were prepared individually in deep-well 96-well plates. A select number of peptides that fell short of the range at no dilution were dried via SpeedVac (200 µL) and reconstituted in 20 µL 0.1% FA. 10 µL of each peptide dilution was pooled together into a final normalized sample. The peak height of each peptide was analyzed as previously stated.

**Details of Proteome Discoverer DDA search**

DDA raw files were searched using Proteome Discoverer version 2.4.0.305. The Processing Workflow tree included the following nodes: Spectrum Files RC, Spectrum Selector, Sequest HT, and Percolator. Raw files from WTC11 or AS192 synthetic peptide experiments were searched against the WTC11 protein database as well as a common contaminants protein database. Raw files form BSA benchmarking experiments were searched against the same contaminants protein database in addition to the Uniprot UP000005640 human proteome with the manual addition of the BSA protein sequence.

Spectrum Files RC settings were: Trypsin (Full), 20 ppm precursor mass tolerance, 0.5 Da fragment mass tolerance, static modifications were +304.207 Da or +313.231 Da on any N-terminus, depending on experiment, and +57.021 Da on C. Regression was set to non-linear regression model and coarse parameter tuning. The Spectrum Selector settings were: use MS1 precursor, use isotope pattern in precursor re-evaluation, provide profile spectra automatically, 350-5000 Da for minimum and maximum precursor, with a minimum peak count of one. Scan Event Filters were: FTMS mass analyzer, MS2 for MS order, HCD activation type, minimum and maximum collision energies set to 0 and 1000 respectively, Full Scan type, and position ion polarity. A peak filter of 1.5 S/N was used. The Replacements for Unrecognized Properties were set to: automatic charge, FTMS mass analyzer, MS2 MS order, HCD activation, positive ion polarity, MS resolution of 60000 and MSn resolution of 30000. The Precursor Pattern Extraction was set to 2.5 Da clipping range before and 5.5 Da clipping range after. Sequest HT was parameterized as follows: fully tryptic enzymatic digestion with a maximum of 2 missed cleavage sites allowed, minimum and maximum peptide lengths were set to 6 and 85, respectively, and monoisotopic precursor and fragment ion mass tolerances were set to 15 ppm and 0.05 Da, respectively. In addition, dynamic modifications allowed were the following: TMTpro (+304.207 Da) on peptide N-termini and K, shTMTpro (+313.231 Da) on peptide N-termini and K, Oxidation (+15.995 Da) on M, Acetyl (+42.011 Da), Met-loss (-131.040 Da), and Met-loss+Acetyl (-89.030) on protein N-terminus. Static modification was Carbamidomethyl (+57.021 Da) on C. A concatenated target/decoy strategy was used with validation based on q-value, and a strict Target FDR value of 0.01 was applied. Peptides were further filtered using the Percolator PEP score of ≤ 0.01 using the Peptide Spectral Match (PSM) results file.

The Consensus Workflow tree nodes were: MSF Files, PSM Grouper, Peptide Validator, Peptide and Protein Filter, Protein Scorer, Protein Grouping, Protein FDR Validator, and Protein Annotation. Spectra storage settings were set to Identified or Quantified and Feature Trace storage was set to All. Merging mode was set to Globally by Search Engine Type and reported FASTA title lines were set to Best Match. A standard title line rule was used. The following PSM Filters were used: Max Delta Cn of 0.05, Max Rank of 0, Max Delta Mass was 15 ppm. A 75 percent site probability threshold was used for Peptide Group Modifications. For the Peptide Validator node, the following settings were used: (General) Automatic Validation with controlling for peptide level error rate if possible, strict PSM and Peptide Target FDR of 0.01, relaxed PSM and Peptide Target FDR of 0.05, (Specific) validation based on q-value, automatic target/decoy selection for PSM level FDR calculation based on score. For the Peptide and Protein Filters, the Peptide Confidence as set to at least High, lower confidence PSMs were not retained, minimum peptide length was 6, peptides without protein reference were not removed. A minimum of 1 peptide per protein was required. Default settings were used for Protein Scorer. For Protein Grouping, strict parsimony principle was applied. For the Protein FDR Validator, the strict and relaxed Target FDR settings were 0.01 and 0.05, respectively. The following Protein Annotation aspects were used: Biological Process, Cellular Component, and Molecular Function.
